# Automated head-fixation training system with high levels of animal participation in psychoacoustic tasks

**DOI:** 10.1101/2022.11.29.518279

**Authors:** Ji Liu, Kate Maximov, Patrick O. Kanold

## Abstract

Many animal training paradigms rely on head-fixation. Head-fixation training is typically laborious and can benefit from automation to relieve the workload as well as to reduce the variability in the training outcome. Several groups have reported successful implementations of such systems, but throughput varied greatly across groups. In addition, most studies relied on brief periods head-fixation sessions (≤ 1 minute) to reduce the potential stress on the animal. Here, we report the design of a new system that could achieve head-fixation sessions on the order of minutes with high participation rate from the animal (100%). Throughout the training period, each mouse performed a total of close to 40 minutes of head-fixation training on average on each day and learned common psychoacoustic tasks, i.e., tone detection and tone discrimination. Our system can achieve highly efficient training with minimum idling time, providing an opportunity for combinations with high-end neural recording equipment to achieve maximum training and data collection efficiency.

## Introduction

One of the fundamental goals of neuroscience is to understand how neuronal activities contribute to the individual’s behavior. To answer this question typically requires monitoring the neurons’ activities while the individual is engaged with certain behavior. Although head mounted neuronal recording devices have seen significant innovations recently (Fan et al., 2011; Resendez et al., 2016; Zong et al., 2017), presenting repeated precisely controlled sensory input during behavior, e.g., gratings on a screen, still requires the restraint of the animal’s head (Guo et al., 2014). Head-fixation training combined with modern electrophysiology and imaging techniques have provided a wealth of new knowledge on how the brain functions to support an assortment of behavior (Aguillon-Rodriguez et al., 2021; Burgess et al., 2017; Long and Lee, 2012; Sofroniew et al., 2014). Nevertheless, behavior training can be laborious and typically requires that the animal progressively moves through different stages of training (Aguillon-Rodriguez *et al*., 2021; Burgess *et al*., 2017; Guo *et al*., 2014). A considerable effort has been made to automate the training process (Bernhard et al., 2020; Francis et al., 2019; Francis and Kanold, 2017; Maor et al., 2020; Winslow et al., 2021), including head-fixation behavioral training combined with automated neurophysiological recordings in behaving animals (Aoki et al., 2017; Hao et al., 2021; Murphy et al., 2016; Murphy et al., 2020; Scott et al., 2013).

Compared to the automated free-moving animal training system (Francis *et al*., 2019; Francis and Kanold, 2017), head-fixation training system offers the advantage of identifying individual animals, typically through RFID (Murphy *et al*., 2016; Murphy *et al*., 2020). However, the head-fixation protocol varied greatly across these studies, and typically each head-fixation episode lasted from several seconds to 1 minute (Hao *et al*., 2021; Murphy *et al*., 2016; Murphy *et al*., 2020; Scott *et al*., 2013) while Aoki et al. adopted a fixed training period of 20 to 30 minutes (2017). Although the average accumulative head-fixation time per day varied from ∼18 min (Murphy *et al*., 2020) to exceeding 1 hour (Hao *et al*., 2021), it did not mimic manual head-fixation training and recording sessions that typically lasted continuously for tens of minutes. Thus, we aimed to design an automated head-fixation system that could mimic manual training as closely as possible and to achieve a higher participation rate from the animal with potentially longer head-fixation episodes.

We based our design on previous publications (Murphy *et al*., 2016; Murphy *et al*., 2020) and we designed custom headplates and a novel head-fixation mechanism and strategy. We also integrated RFID identification into our system, allowing us to customize the training for each mouse. The improved setup allowed us to achieve 100% participation rate out of 14 mice (4 litters) tested in this study. On average, each mouse participated in voluntary head-fixation training for a total close to 40 minutes per day, while the average duration per head-fixation episode approached 15 minutes as training progressed. Therefore, the design of the system presents an opportunity to further improve the throughput of the automated head-fixation training systems, which enable the training of large cohorts of mice for downstream neurophysiological experiments.

## Methods

### Animals

All protocols and procedures are approved by the Johns Hopkins Institutional Care and Use Committee. We used 7 male and 7 female adult mice that were the F1 generation of Thy1-GCaMP6s (JAX# 024275) crossed with CBA/CaJ mice (JAX# 000654). These 14 mice came from 4 litters (3 males, 4 males, 3 females, 4 females). The F1 generation has minimum hearing loss throughout their lifespan (Bowen et al., 2020; Frisina et al., 2011; Shilling-Scrivo et al., 2021). The mice used in this study ranged from 3 to 6 months old when the training started.

### Automated training apparatus

Following a similar approach to previous studies (Murphy *et al*., 2016; Murphy *et al*., 2020), the main component of our automated homecage training system consisted of a tunnel which the animal entered for head-fixation training (Figure 1A). Together with the custom designed headplate mounted onto the skull of the mice, the tunnel guided the mice to reach the end where the head-fixation was performed. The custom headplates were 3D printed in aluminum. Specifically, the custom headplate had two guide rods extending laterally (Figure 1B). On the other hand, the cross-section of the tunnel consisted of a trapezoid area in the center that accommodated the body of the mice and an adjoining rectangular area on the top that represented the slots where the guide rods moved along while the mice entered the tunnel (Figure 1B, bottom). The slots gradually shrink in the vertical direction towards the end of the tunnel to provide more constraint on the head movement of the mice. When the animal reaches the end of the tunnel, an anchor bar driven by a servo pressed the back of the headplate such that both guide rods were pressed against the front wall of the tunnel where a small protrusion further guided the positioning of these rods (Figure 1B, C). The firm push by the anchor bar thus ensured the secure head-fixation. We chose the cylindrical profile of the guide rods as well as their rounded end to allow the animal to have some degree of pitch and yaw movement before reaching the end of the tunnel and fully head-fixated, which encouraged the mice to explore the tunnel in the initial stages of training and helped to reduce their stress and the chance of fear association.

**Figure 1.**
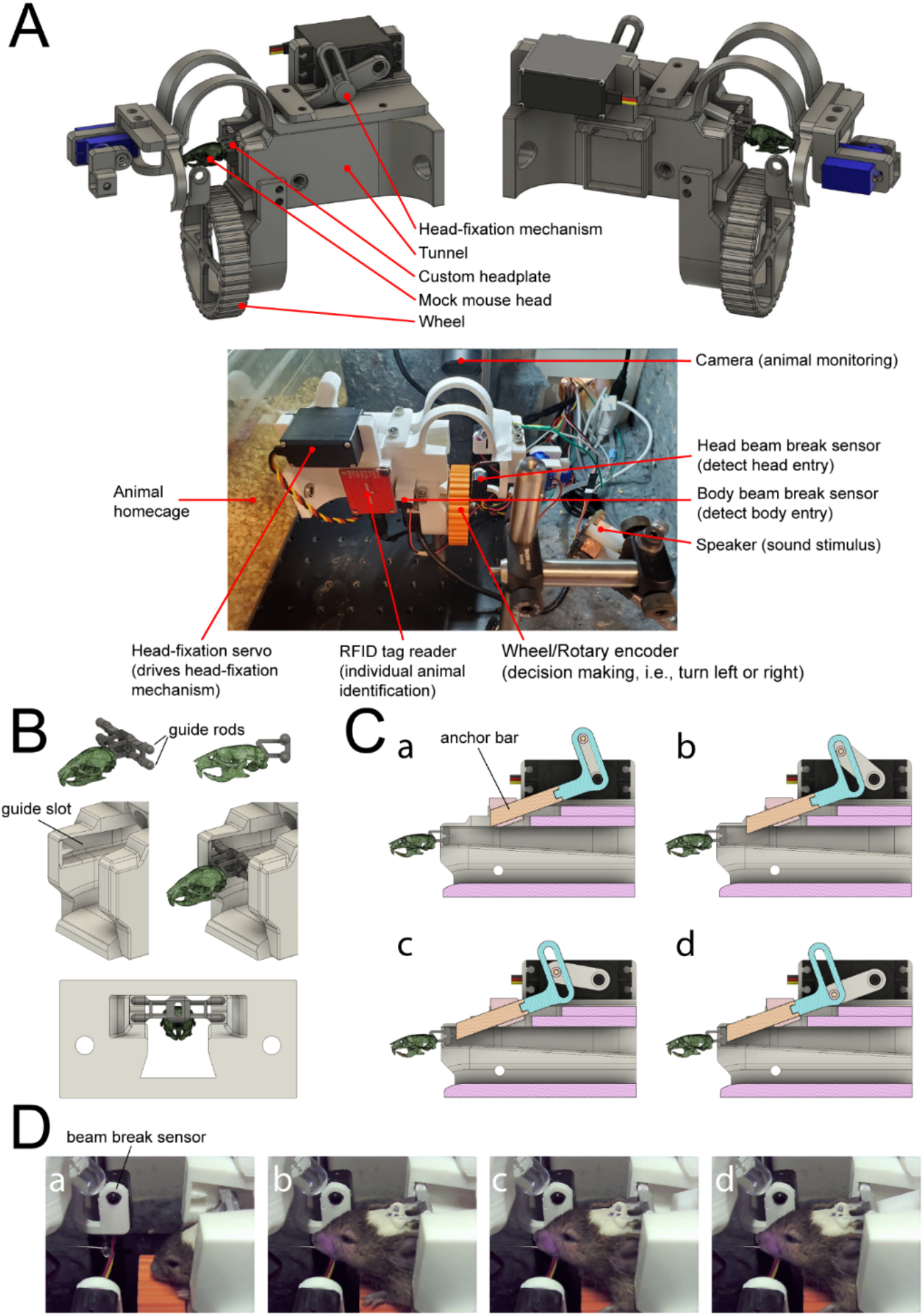
Design of the automated head-fixation training system. (A) Top: two views of the Computer-Aided Design (CAD) of the automated head-fixation system. The system mainly consisted of a tunnel into which the mouse entered for head-fixation training. The tunnel worked in tandem with a custom headplate attached to the back of the skull of the mouse. Bottom: photo of the actual system. The tunnel also housed various electronics, including the head-fixation servo, the RFID tag reader for identifying individual mouse, the wheel attached to a rotary encoder for reporting behavior, the beam break sensors for detecting the mouse’s body and head entries, the speaker for sound delivery and the camera for mouse monitoring. (B) Top: two views of the custom headplate relative to the mouse’s skull. The most important feature of the headplate is the two guide rods placed vertically. Middle: two views of the end of the tunnel with and without the mouse’s head entry. The guide rods of the headplate fit in the two slots on the lateral side of the tunnel. A small protrusion that fit between the guide rods further guide the headplate into the final position. Bottom, the rear view of the tunnel and the headplate in the head-fixation position. The two slots on the lateral side of the tunnel shrank in size as the mouse approach the head-fixation position. (C) Panels (a)-(d) show the head-fixation mechanism. The cross-sections of the tunnel were shown. The mechanism translates the rotation motion of the servo into the linear motion of the anchor bar. When fully extended, the front face of the anchor bar pushed against the back of the headplate, preventing the exit of the mouse’s head. (D) Panels (a)-(d) show the sequence of actual mouse head-fixation. The mouse entered to consume the entry reward while triggering the beam break sensor, which in turn activated the head-fixation servo that moved the anchor bar forward to achieve head-fixation.

The main mechanism for the head-fixation is the servo-driven anchor bar (Figure 1C). We designed a simple mechanism that translated the rotation motion of the servo into the linear motion of the anchor bar. The anchor bar moved along a direction with a tilt of 20 degrees with respect to the horizontal plane while the front face of the anchor bar had its normal direction aligned with the horizontal plane. Under the default configuration, the anchor bar was fully retracted and did not hinder animal entering the tunnel. When fully extended, the front face of the anchor bar pressed onto the back of the headplate to achieve head-fixation.

To detect the entry of the mouse, we used two pairs of infra-red beam break sensors (Adafruit, product ID 2168). One pair was placed around the middle of the tunnel to detect the body entry of the mouse, while the other pair was placed at the end of the tunnel to detect the mouse’s head entry (Figure 1A). A digital camera (Amazon, USB-USBFHD01M-SFV(5-50mm)) was set up to monitor the mouse’s actions within the tunnel. Once the mouse activated the body entry beam break sensor, the camera started recording until the mouse left the tunnel. At the same time, the RFID reader (Sparkfun ID-20LA) placed at the right-hand side of the tunnel would read the RFID tag implanted beneath the skin of the mouse near the right abdomen. The training stage as well as the head-fixation time depended on the mouse’s ID. The program only proceeded to head-fix the mouse once the ID was determined successfully from the RFID reader. After that, the mouse needed to fully enter the tunnel and activate the head entry beam break sensor with its snout before the anchor bar began extending towards the back of the headplate. The speed of the anchor bar movement was dependent on how far the front face of the anchor bar was away from the headplate. As the anchor bar approached the headplate, the speed would gradually decrease to the minimum speed to reduce the impact force onto the headplate. This feature helped to reduce the potential stress the mouse could experience. It took around 1.5 to 2 seconds for the anchor bar to reach the fully extended position for head-fixation and if the mouse failed to maintain its head position during this period, the anchor would begin retracting. We positioned the head entry bream break sensor such that to activate it not only required the mouse’s full head entry, but also that the mouse held its head in a relatively horizontal position. Because of the cylindrical profile of the guide rods, the mouse could easily tilt the head down about the axis provided by the guide rods and deactivate the sensor. Thus, these design features required that the mouse voluntarily meet the conditions for head-fixation, i.e., holding the head in the proper position for a sufficient amount of time. In addition, the sound of the servo movement also clearly signaled to the mouse the initiation of the head-fixation process. Unless the mouse was willing to perform head-fixation voluntarily, it could use the sound cue to abort the process. The ease with which the mouse could abort head-fixation as well as the soft impact of the anchor bar onto the headplate helped to reduce the overall stress of the mouse during the head-fixation process and promoted the mouse’s willingness to perform head-fixation.

Although head-fixation required that the mouse’s head be held in a relatively horizontal position, this condition can be typically met even if the animal’s head was slightly tilted down, i.e., only the upper guide rod was in contact with the front wall of the tunnel (a perfectly horizontal position requires that both guide rods are in contact with the front wall, as in the final head-fixation position). The extending anchor bar thus came in contact first with the lower half of the back of the headplate and pushed it to rotate about the axis defined by the upper guide rod until both upper and lower guide rods were pressed firmly against the front wall. From our observation, this design allowed robust head-fixation results even if the mouse’s head was not in an optimal position. The rotating axes provided by the guide rods made the corrections of the head’s position straightforward. Together, we believe our head-fixation design greatly reduced the stress the mouse could experience, which is also manifested in their heightened engagement in performing voluntary head-fixation training.

At the end of the tunnel, the mouse collected water reward by licking a feeding needle that served as the waterspout. The waterspout was controlled by a servo such that the waterspout could move out of the reach of the tongue of the mouse. This design helped to reduce uninstructed licking activities during tone presentations by retracting the waterspout.

The auto head-fixation mechanisms were controlled by a custom written LabView program, which also communicated with an Arduino that controlled the servo movements, i.e., the head-fixation servo and the waterspout servo.

### Voluntary head-fixation training

To train mice to perform voluntary head-fixation training, we designed 4 progressive training stages (HF1 to HF4) that gradually familiarized mice with the tunnel and accustomed them to head-fixation. In the first stage (HF1), the goal is to encourage the mouse to enter the tunnel and to associate this behavior with collecting water reward. A mock tunnel was used that did not have the other electronic components, e.g., head-fixation servo. Instead, a custom designed plate was mounted on the top of the tunnel that had four different positions to insert the waterspout. The waterspout was the same as the one used in normal animal housing. On consecutive days of training, the waterspout was moved progressively by one slot from the initial position closest to the homecage to the final position at the end of tunnel. This process typically required 4 to 5 days. Before the first day of training in this stage, mice were first water deprived for 24 hrs. On every day, mice were first weighed and then introduced to the cage where the mock tunnel was attached for 2 hours, and the mice received no other water source. The mice’s activities were monitored using a digital camera to verify that they indeed entered the tunnel to drink water. After the session concluded, we weighed the mice again to confirm that they consumed water. For any mouse that did not enter the tunnel to consume water, the waterspout would be moved closer to the cage by one slot to further accustom them to the tunnel, until they robustly enter the tunnel for water reward. Mice quickly learnt to enter the tunnel for water consumption, after which they were transitioned to stage HF2.

In the second stage (HF2), the mice were introduced to the homecage where the tunnel with the full suite of head-fixation mechanisms and electronics was installed. The mice were required to learn to break the head entry beam break sensor in order to collect water reward. However, no head-fixation was introduced in the stage. Most importantly, this stage accustomed the mice to hold their heads in the proper position for future head-fixation. The subsequent events that happened when the mouse entered the tunnel were the following. When the mouse activated the body entry beam break sensor, one drop of water was dispensed (5-7 μl) at the waterspout to attract the mouse to enter further. Once the mouse reached the end of the tunnel and activated the head entry beam break sensor, one water drop was dispensed every 2 sec for as long as the beam break signal remained high but no more than 5 drops. Thus, to collect all 5 drops of water reward, the mouse would need to hold the head still for around 10 sec. If the mouse failed to maintain the head position and deactivated the beam break sensor or all 5 drops of water were dispensed, the water drop counter reset and the mouse would need to re-initiate the process. However, within one entry, the mouse was free to initiate multiple head entries to collect as many drops of water reward as it desired. This stage allowed the mice to be further accustomed to the head-fixation setup and the mice robustly performed head entries to collect water reward. This stage lasted 2 to 3 days. As in the first stage, we weighed the mice both before and after the training to ensure that the mice collected water reward, i.e., their body weights increased after training.

In the third stage (HF3), actual head-fixation was introduced and accompanied the head entry water reward collection. This stage was similar to the second stage except that when the mouse performed head entries, the dispense of water reward depended not on activating head entry beam break sensor, but on the completion of head-fixation. The anchor bar start extending once the mouse’s head activated the beam break sensor and the movement was considered complete once the anchor bar was in the fully extended position. Only then were the 5 drops of water started dispensing with an interval of 2 seconds. Thus, the mouse was head-fixed for around 10 seconds for each of these mini head-fixation sessions. Similar to the second stage, the mouse was free to initiate as many head-fixation sessions as desired within one tunnel entry. Unlike previous studies that initially used loose head-fixation to prevent mice from producing fear association (Murphy *et al*., 2016; Murphy *et al*., 2020), we started by exposing the mice with full-strength head-fixation with a short duration and we did not notice any adverse effect. In fact, all mice performed multiple head-fixations on the first day of this stage, suggesting that this process produced very little stress for them. This was likely further aided by the fact that the final head position for head-fixation was similar to that of the last stage, and that the mice have been well accustomed, and our head-fixation mechanism produced little discomfort. This stage lasted another 3 days.

In the fourth stage (HF4), we replaced the mini head-fixation session in stage 3 with the actual behavioral task. Thus, except for the entry reward dispensed when the mouse entered the tunnel, all other water rewards were collected during the behavioral task. The goal of this stage is to produce behavioral training sessions similar in period to that of our manual training sessions (∼30 min). However, so far the mouse had only been exposed to short periods of head-fixations, we thus start the stage with short training sessions, with minimum duration and maximum duration both equal to 0.5 min. We gradually increased the minimum and maximum session duration according to the following rule: for every 5 head-fixation sessions, the minimum duration increased by 0.25 min while the maximum duration increased by 0.5 min. Thus, as the mouse was more exposed to the head-fixation trainings, the longer they were required to perform. However, the minimum duration cannot exceed 3 min while the maximum duration cannot exceed 30 min. For a head-fixation training session to end, the minimum training duration had to be reached, after which the session would end if the mouse failed to make any hits in consecutive 5 trials or the time reached the maximum duration. After the session concluded, the anchor bar would fully retract and wait for the mouse to retract its snout. However, after the snout retraction, the mouse was free to choose to fully exit the tunnel or to initiate another head-fixation training session. The mice stayed in this stage while making progress with respect to the behavioral task.

In all of the above 4 stages, the mice were introduced to the head-fixation setup for a limited amount of time per day rather than given unlimited access. In stage two through stage four, the total amount of time one cage of mice was exposed to the training setup per day equaled to 40 min times the number of mice in the cage. We chose this time limit for several reasons. First, it is similar to the time limit of our manual training paradigm. Second, limited access to the training setup helped to regulate the motivation level of the mice and encouraged them to perform better given the limited time to perform the task. Third, this arrangement can allow the same head-fixation setup to be used by multiple cages of mice, where each cage consisted of 3 to 4 mice, further increasing the training throughput. For the above reasons, we advocate limiting access to the training setup even with automated training equipment.

### Tone discrimination task

Using the automated head-fixation training system, we then trained the mice to perform a frequency discrimination task. In the default task, the mice were presented tones between 10 and 40 kHz and the mice were required to turn a wheel placed in front of them either to their right in response to low frequency tones (<20 kHz) or to the left in response to high frequency tones (>20 kHz). In a subset of randomly chosen mice, the turn directions were reversed with respect to the tone frequencies. Specifically, the tones were sinusoidally amplitude modulated with 10 Hz and the maximum amplitude was calibrated to 70 dB SPL. Each tone was 2 seconds in duration. However, after a grace period passed since the tone onset (0.1 sec) where any wheel turning behavior was discounted, the mouse was free to make a choice by turning the wheel in either direction, up until 4 seconds after the tone onset. A choice was considered made by the mouse whenever it turned the wheel beyond a certain threshold (5 or 10 degrees, depending on the training stage) in either direction. Whenever a choice was made, the feedback was immediately conveyed to the mouse either by presenting the water reward in a hit trial, i.e., the retractable waterspout would tilt up and dispense one drop of water (5-7 μl), or by presenting a silent timeout of 4 to 8 sec in trials with incorrect choices. In more advanced training stages (stage 2 to stage 5), tactile feedbacks were also presented by activating a vibration motor attached to the side of the auto head-fixation tunnel for 0.2 sec in incorrect trials. What’s more, if the behavioral choices were made before the tone offset, the tone presentation was also immediately stopped, and the feedback was presented. Thus, the trials were structured to convey the consequence of the mice’s behavioral choices as saliently as possible.

We trained the mice through several stages that differed in the stimuli used and other conditions. We constructed the mice’s psychometric curves from the last stage where all tones were presented. The tones we presented were of the following frequencies: 10, 14.1, 16.8, 18.3, 19.1, 20, 20.9, 21.8, 23.8, 28.3, 40 kHz. These tones were symmetric around 20 kHz in the logarithmic space and corresponded to the following octaves in reference to 20 kHz: -1, -0.5, -0.25, -0.125, -0.0625, 0, 0.0625, 0.125, 0.25, 0.5, 1. Thus, these tones were most densely sampled around 20 kHz, the sensory boundary. Table 1 listed the frequencies of tones used in each stage along with the probability they were presented with. In stage D1 and D2, the mice were presented with tones that were furthest away from the 20 kHz, and subsequent stages gradually added more tones close to 20 kHz. In stage D1 and D2, to correct any turn direction bias, we implemented a simple bias correction mechanism that kept presenting the same tone until the mouse performed correctly. However, after a hit trial, all frequencies were selected randomly. In stage D1 and D2, the mouse needed to perform with at least 80% hit rate within any consecutive 50 trials to advance to the next stage. After stage D2, stage advancement was no longer conditioned on the mouse’s performance. Rather, the mice simply needed to perform enough trials within each stage D3 and D4 to advance to the next stage. In stage 5, all tone frequencies were presented, including 20 kHz. Turning in either direction in response to 20 kHz resulted in water reward. In stage D3, D4 and D5, all tones except for 10, 14.1, 28.3, 40 kHz were presented with a probability of 0.05, while the presenting probability for these 4 tones were calculated accordingly. The tone discrimination psychometric curve was constructed from the data collected in stage 5. The tones at the two ends of the spectrum were presented more often to ensure a reasonably high hit rate to engage the mice in the task, while the tones in the middle of the spectrum were considered harder and only presented to measure the psychometric curve.

**Table 1.** Frequency discrimination task stage parameters.

|  | **stage D1** | **stage D2** | **stage D3** | **stage D4** | **stage D5** |
| --- | --- | --- | --- | --- | --- |
| **Tone frequency (kHz)** | 10, 40 | 10, 14.1, 28.3, 40 | 10, 14.1, 16.8, 23.8, 28.3, 40 | 10, 14.1, 16.8, 18.3, 21.8, 23.8, 28.3, 40 | 10, 14.1, 16.8, 18.3, 19.1, 20, 20.9, 21.8, 23.8, 28.3, 40 |
| **Presenting probability** | 0.5, 0.5 | 0.25, 0.25, 0.25, 0.25 | 0.225, 0.225, 0.05, 0.05, 0.225, 0.225 | 0.2, 0.2, 0.05, 0.05, 0.05, 0.05, 0.05, 0.2, 0.2 | 0.1625, 0.1625, 0.05, 0.05, 0.05, 0.05, 0.05, 0.05, 0.05, 0.1625, 0.1625 |
| **Bias correction** | Y | Y | N | N | N |
| **Stage advancement criterion** | Hit rate ≥ 0.8 for any consecutive 50 trials in one session | Hit rate ≥ 0.8 for any consecutive 50 trials in one session | Performed ≥ 50 trials for at least 5 of the past 10 sessions | Performed ≥ 50 trials for at least 5 of the past 10 sessions | N/A |
| **Decision threshold (deg)** | 5 | 10 | 10 | 10 | 10 |
| **Timeout (sec)** | 4 | 4 | 6 | 8 | 8 |
| **Tactile feedback duration for incorrect trials (sec)** | 0 | 0.2 | 0.2 | 0.2 | 0.2 |

During training, the sound waveform was generated by NI-USB6215 (National instrument) and routed to an ED1 speaker driver (Tucker-Davis Technologies), which drove an ES1 open field speaker (Tucker-Davis Technologies) for playout. The tones were calibrated with a B&K microphone (Bruel & Kjaer 4944-A).

### Tone detection task

In a subset of mice, we also trained them to perform a tone detection task after they mastered the tone discrimination task. The frequencies of the tones were equally sampled in the logarithmic space from 7.1 to 56.6 kHz with a density of 5 tones per octave (16 tones in total).

The tones were not amplitude modulated and were of 0.2 sec in duration. Each trial had a maximum duration of 0.5 sec since the tone onset and the mouse needed to turn the wheel in either direction before the end of the trial to collect water reward. No punishment in terms of timeout or tactile feedback was given in miss trials where the mouse failed to respond to the tone within the 0.5-sec trial period. We introduced the mice to 3 stages of training, where in each stage all tone frequencies were presented but the sound levels varied (stage T1 through T3, Table 2). To advance beyond stage T1 required the mouse to detect 80% of tones within any consecutive 50 trials, while to advance beyond stage T2 only required performing sufficient number of trials (Table 2). We constructed the behavioral audiogram using data collected in stage 3.

**Table 2.** Tone detection task stage parameters.

|  | **stage T1** | **stage T2** | **stage T3** |
| --- | --- | --- | --- |
| **Sound level (dB SPL)** | 70 | 70, 60, 50, 40 | 70, 60, 50, 40, 30, 20, 10 |
| **Decision threshold (deg)** | 5 | 15 | 15 |
| **Stage advancement criterion** | Hit rate $\geq 0.8$ for any consecutive 50 trials in one session | Performed $\geq 50$ trials for at least 5 of the past 10 sessions | N/A |

### Surgery

To prepare the mice for automated head-fixation training, custom designed headplates were mounted to their skull. The surgery was performed under anesthesia using isoflurane (4% induction, 1.5% maintenance). First, the hair on the top of the head was removed. Next, the skin as well as the soft tissues beneath were removed with disinfected scissors and scalpel blades. We next exposed the dorsal and the caudal sides of the skull. Then we made repeated measurements of the z-axis reading of the lambda and bregma landmark on the skull and made adjustments such that the readings of the two landmarks were no more than 200 μm. Next, we attached the headplate to the stereotaxis manipulator through a 3D printed adapter and positioned the headplate such that both the midline of the headplate and the midline of the head were aligned and the front of the headplate was aligned with the Lambdoid sutures. Next, we applied dental cement to secure the headplate onto the skull (C&B Metabond). Finally, a small incision was made over the skin coving the right abdomen and the RFID tag was inserted before suturing the skin. After the surgery, the mouse was placed under a heat lamp for recovery for 30 min before being placed back in the homecage. Training began 5 to 7 days after the surgery was performed.

## Results

### Design of the automated head-fixation training system

We aimed to design an automated head-fixation training setup with potentially improved task engagement from the animals that rivaled manual head-fixation training. To achieve this goal, we designed a new custom headplate and a corresponding new head-fixation mechanism. The new headplate and mechanism allowed gradual habituation to the tunnel and head-fixation mechanism. Together with the training protocol that systematically guided the mice through different training stages, we were able to significantly improve the time each mouse spent performing voluntary head-fixations.

Based on previous studies (Murphy *et al*., 2016; Murphy *et al*., 2020), our design mainly consisted of a tunnel that was attached to the housing cage of the mice (Figure 1A). During training, individual mouse entered the tunnel and at the end of the tunnel the controlling software detected the mouse’s head entry through beam break sensors. The head-fixation mechanism was then engaged to hold the mouse’s head in position, and the mouse performed voluntary head-fixation training. The tunnel was equipped with an RFID reader such that it could identify individual mouse through the RFID tag embedded in beneath the skin. Two pairs of beam break sensors were employed to better determine the position of the body and the head of the mouse. Our system used a wheel placed in front of the mouse as the instrument for reporting choice. The mice indicated their choice by turning either towards their left or right while the wheel movement was readout by a rotary encoder.

To achieve robust voluntary head-fixation behavior, we designed a custom headplate (Figure 1B). The main feature of the headplate was two guide rods that were positioned vertically (Figure 1B). As the animal entered the tunnel, the guide rods fit in the designated guide slots in the tunnel, which gradually shrank in size as the animal approached the end of the tunnel (Figure 1B). In the final head-fixed position, the front of the two guide rods were in contact with the front wall of the tunnel while the back of the headplate was pushed by an anchor bar that prevented the animal from exiting (Figure 1C, D). The rounded profile of the guide rods reduced the contact surface with the guide slots in the tunnel and allowed the mouse to smoothly enter and exit the tunnel. Furthermore, the two axes provided by the two guide rods allowed the mouse to tilt their head up and down (pitch) and the rounded ends of the guide rods allowed the mouse’s head to rotate sideways (yaw). These movements were possible even when they reached the end of tunnel. Thus, without the engagement of the head-fixation mechanism, the mouse could rather freely explore the tunnel without feeling constrained, which in turn encouraged them to enter. This was especially important during the initial habituation stages (stage HF1 and HF2, see Methods) as the mouse could easily enter and exit the tunnel and associate the experience most strongly with collecting water reward, rather than with being “stuck” or being head-fixed.

To transition the mice smoothly to voluntary head-fixation, we designed 4 stages of training to accustom mice to (stage HF1 to HF4, see Methods). Briefly, stage HF1 used a mock tunnel to teach mice to gather water reward from the end of the tunnel. Stage HF2 transitioned the mice to the actual automated head-fixation tunnel while HF3 introduced mini head-fixation sessions of 10 sec in duration. The actual behavioral task was introduced in stage HF4. By stage HF4, the mice showed robust voluntary head-fixation behavior with no obvious signs of stress We trained 4 cages of mice (n=14) in this study. On each day, each cage of mice was introduced to the automated head-fixation system for a restricted amount of time (40 minutes times the number of mice in the cage, see Methods). All 14 mice successfully went through stage HF1 to HF4, showing that our automated head-fixation system as well as our training protocol induced little stress in our animals.

### Automated head-fixation system achieves high animal engagement and stable training result in a frequency discrimination task

In stage HF4, we first trained the mice to perform a frequency discrimination task, which required the mice to turn a wheel placed in front of them rightward if low frequency tones were presented (<20 kHz) or to turn leftward when high frequency tones were presented (>20 kHz, Figure 2A). The lowest and highest tones used in this task were of 10 and 40 kHz respectively. The frequency discrimination task also consisted of different stages where intermediate tones close to 20 kHz were progressively added to the stimuli set (see Methods).

**Figure 2.**
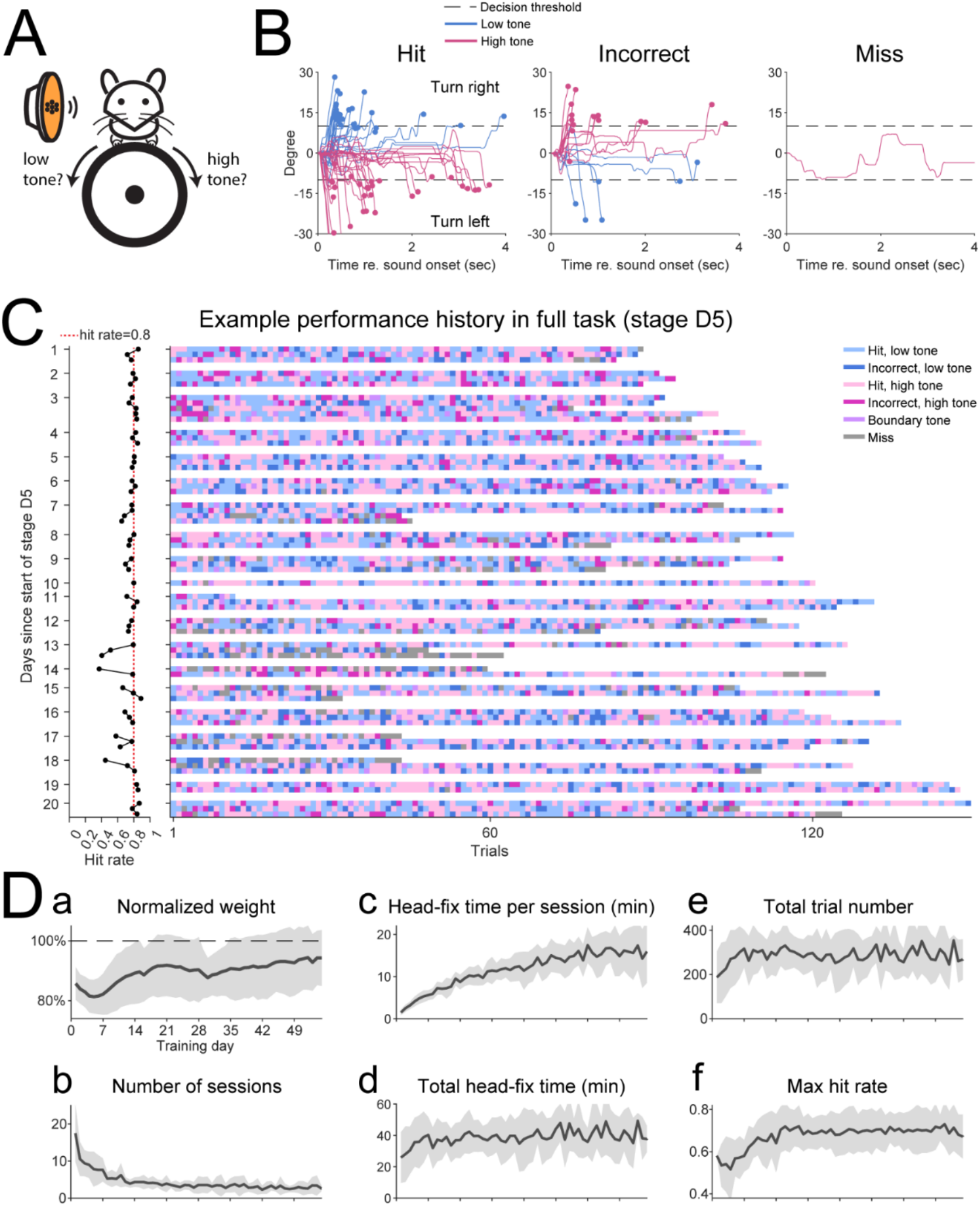
The automated head-fixation training system achieved high animal engagement and stable animal performance. (A) Tone discrimination paradigm: the mouse was trained to turn the wheel towards its right when a low frequency tone was presented (<20 kHz) and to turn the wheel towards its left when a high frequency tone was presented (>20 kHz). Turning in either direction was rewarded when 20 kHz was presented. (B) The wheel movement from one example session in training stage D2 as a function trial outcome was shown. The wheel movement that crossed the decision threshold was considered an actual decision. In miss trials the wheel movement failed to cross this threshold. (C) The performance history of one mouse in the full discrimination task (final training stage D5). Day 1 corresponded the first day entering stage D5. Left: the overall hit rate of all sessions performed on each day was plotted. Each dot corresponded to one session. All dots corresponding to the sessions on the same day were connected. The vertical dotted red line indicates a hit rate of 0.8. Right: the trial outcomes of each session were shown color-coded. Each pixel corresponded to one trial while each row corresponded to one session. This example mouse showed stable performance over 20 days of training. (D) The training history of all mice in this study (n=14). (a)-(f) respectively showed the normalized weight, number of sessions performed, head-fixation duration per session, total head-fixation time, total trial number, max hit rate over any consecutive 50 trials as a function of training days. The solid lines represent the mean and the shaded regions represent the standard deviation across mice.

Mice successfully learnt to perform this frequency discrimination task under our automated head-fixation training system. Figure 2B showed an example mouse’s behavior in one session at training stage D2 where 2 low tones (10 and 14.1 kHz) and 2 high tones (28.3 and 40 kHz) were presented. The mouse turned wheel in the correct direction with an overall hit rate of 73% while the peak hit rate over any consecutive 50 trials was 80%. Figure 2C shows the performance history of another mouse in stage D4 where all intermediate tones were presented over 20 days. This mouse had a stable performance with a hit rate around 80% and engaged in multiple head-fixation training sessions per day. These examples show that our auto head-fixation training system could train mouse to perform robustly and reliably.

Figure 2D shows the summary training history for the frequency discrimination task for all 14 mice used in this study. All of the 14 mice engaged in voluntary head-fixation training and went through all stages of the frequency discrimination task. These mice remained healthy throughout the training period, as the mice’s weight remained constant around 90% of their original body weight despite an initial decrease in the first week following the onset of the training (Figure 2Da). This result suggests that mice were able to maintain a healthy level of water deprivation while entirely dependent on the automated head-fixation training system for water. As the training progressed, we observed a decrease in the number of head-fixation training sessions per day (Figure 2Db), which was accompanied by a gradual increase in the duration of each head-fixation session (Figure 2Dc). The product of these two factors, i.e., the total head-fixation duration per day showed an increase over the first week of training while remained constant throughout the rest of the training period (Figure 2Dd). For each cage of mice on a given day, the total time the mice were allowed access to the training system equaled to 40 min times the number of mice in that cage (see Methods). The fact that the average total head-fixation time for each mouse approached this time limit (40 min) suggests that mice fully utilized their access to the training system and the idle time of the system was minimum. This result further suggests that the mice were highly engaged and likely experienced little stress from the head-fixation system and that our training system operated in a highly time efficient regime. Similar to the total head-fixation time, the total number of trials performed on each day remained constant around 300 after an initial increase in the first week, suggesting that the two factors linearly scale with each other (Figure 2De). Finally, the max hit rate over any consecutive 50 trials within a session in a given day (Figure 2Df) showed a steady increase over the course of 3 weeks following the training onset and stabilized around 70%. Together, these results show that our automated head-fixation training system can achieve robust and stable training result. Most importantly, our design offered great improvement in terms of the animals’ participation rate and their engagement.

Our automated training system was able to identify individual mouse by reading their embedded RFID tag, which allowed us to customize training for each mouse. In the default frequency discrimination task, low frequency tones were mapped to turning right while high frequency tones were mapped to turning left. Figure 3A shows the psychometric curves obtained from 10 mice trained with the default task. All but 2 mice showed reasonable performance while these 2 mice showed significant bias towards turning left or right (Figure 3A, orange and purple curves). In a subset of 4 mice, we reversed this relationship, and the mice learnt the reversed mapping between frequency and the turn direction (Figure 3B). These mice trained alongside with other mice with the normal turn direction setting, which was made possible by the reliable identification of individual mouse through their RFID tags. Figure 3C shows the average psychometric curve obtained from all but the 2 mice with a significant turning direction bias. The mean psychometric curve was well described by a sigmoid function.

**Figure 3.**
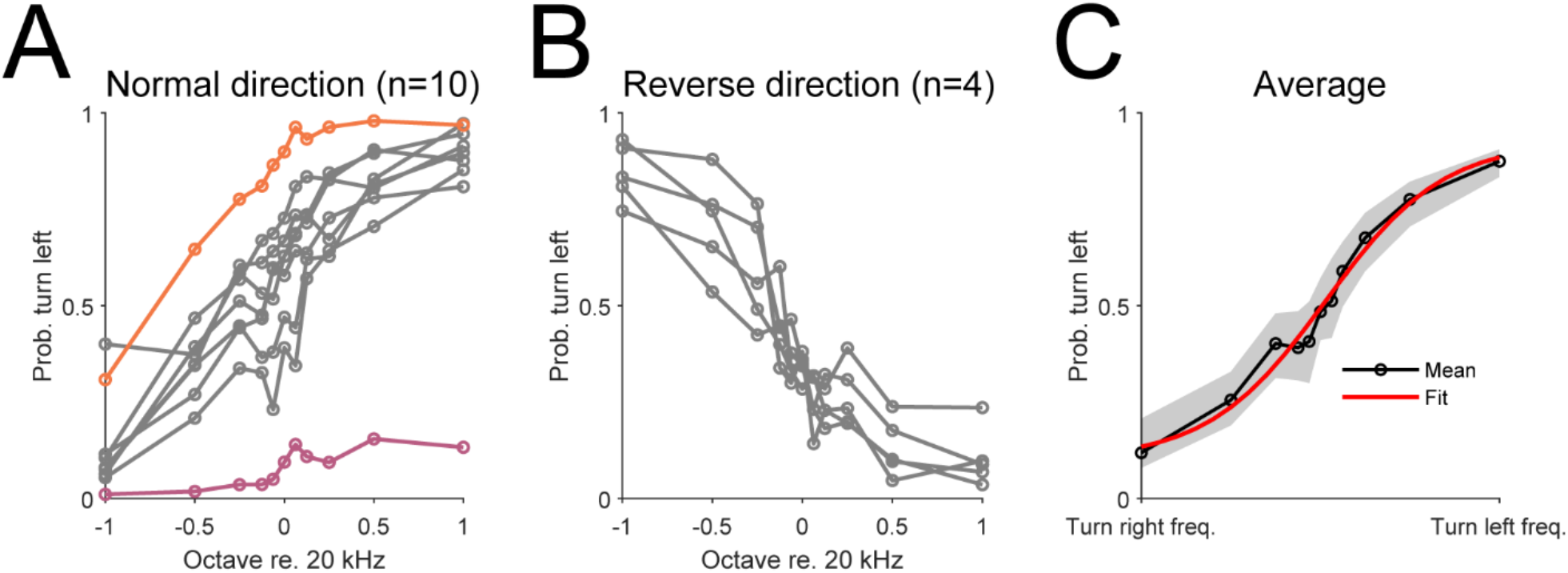
Psychometric curves of mice trained under the automated head-fixation system. (A) Psychometric curves of individual mouse trained in the default frequency discrimination task (n=10). All but 2 mice (orange and purple curves) showed reasonable performance. (B) The psychometric curves of 4 mice trained with the reversed frequency-action mapping. (C) The average psychometric curves of 12 mice (excluding the mice with significant left or right turn biases, i.e., orange and purple curves in (A)). The red curve represents a sigmoid function fit.

Together, these results suggest that our training system was well suited for individualized psychoacoustic experiments and was capable of measuring psychoacoustic abilities of individual mice. Moreover, our system is able to customize the training based on the behavioral performance of each mouse.

### Automated head-fixation system is capable of training mice to perform tone detection tasks

Aside from discrimination tasks, detection tasks are also commonly used psychoacoustic paradigms. As a further proof of concept, we trained a subset of the mice (1 cage of 4 mice) to perform a tone detection task after they have learnt the discrimination task and we obtained their psychometric curves (Figure 4A). This task consisted of 16 tones spaced equally in the logarithmic space from 7.1 to 56.6 kHz and the sound level varied from 10 to 70 dB SPL with a step of 10 dB SPL. The tones were 0.2 sec in duration and in each trial the mouse had 0.5 sec from the onset of the tone to respond. Figure 4B shows the wheel movement traces by one mouse from an example session, where a hit was achieved if the wheel movement passed the decision threshold by the trial’s end or the trial was considered a miss. The example shows that the mouse can robustly report the presence of the tone by turning the wheel. Figure 4C shows the detection rate as a function of tone frequency and sound level, plotted separately for individual mouse and the group average. Typically, the hit rate increased as the sound level increased and showed frequency dependency. We constructed the average behavioral audiogram by interpolating the sound level with a hit rate of ∼40% at each sound frequency (Figure 4C, right). Our behavioral audiogram showed the maximum sensitivity 14.1 kHz, consistent with results reported by previous studies (Francis *et al*., 2019; Koay et al., 2002; Radziwon et al., 2009). We further examined the hit latency, defined as the time the wheel movement passed the decision threshold, as a function of frequency and sound level (Figure 4D). Qualitatively, the hit latency was the highest among low sound levels and tended to decrease as the sound level increased. Indeed, the hit latency was significantly negatively correlated with the sound level (Figure 4D, right). This result suggests that the softer the sound, the more slowly the mouse responded, which is consistent with finding that the latency of auditory responses increased as the sound level decreased (Klug et al., 2000). Overall, these results show that our automated head-fixation training system was also capable of training mouse to perform tone detection tasks. Together with the frequency discrimination task, our system demonstrated the capabilities to perform conventional psychoacoustic experiments with improved animal engagement and little human intervention.

**Figure 4.**
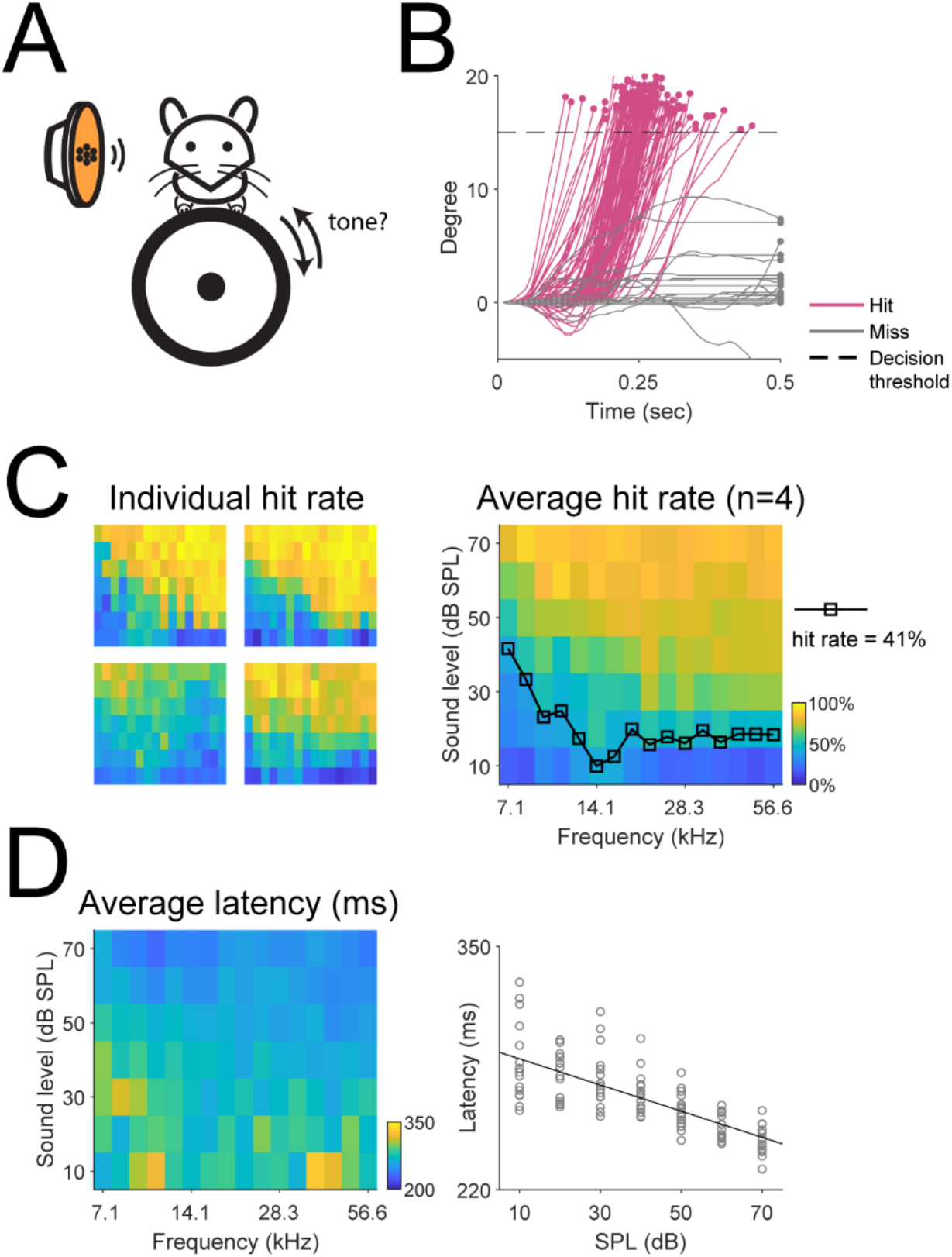
The automated head-fixation training system readily trained mice to perform a tone detection task. (A) We trained a subset of mice (n=4) to perform a tone detection task, where the mice were required to turn the wheel in either direction in response to a brief tone (0.2 sec in duration). (B) The wheel movement traces from an example tone detection session, plotted as a function of trial outcome. (C) Left: the hit rate as a function of frequency and sound level, plotted separately for individual mouse. Right: the average hit rate as a function of frequency and sound level. We used a hit rate of 41% to define the behavioral audiogram. (D) Left: the average hit latency, i.e., the time at which the wheel movement crossed the decision threshold, was plotted as a function of frequency and sound level. Right: the average hit latency was plotted against the sound level. A significant negative relationship exited between the two (slope = -0.70, p=1.51×10^−20^).

## Discussion

In this study we presented an automated head-fixation system that achieved a high animal participation rate and engagement. We trained mice to perform a simple tone detection task as well as a more complex 2AFC discrimination task and we successfully obtained psychoacoustic measurements from individual mice. Furthermore, our system’s ability to reliably identify individual mice allowed them to progress through task stages independently of each other and also allowed us to customize the task parameters for each mouse, e.g., the mapping between frequency and the wheel turning direction. We believe that our system presents an opportunity to further boost the throughput of the automated head-fixation training system.

Head-fixation training can be stressful for the animal and to prevent buildup of stress previous studies typically employed short bouts of head-fixation (less than or equal to 1 minute) (Hao *et al*., 2021; Murphy *et al*., 2016; Murphy *et al*., 2020; Scott *et al*., 2013) while Aoki et al. used a fixed and longer training period of 10 to 20 minutes (2017). Our design of the system operated in the intermediate regime where the minimum duration for each head-fixation session was on the order of minutes (3 minutes eventually). As the training progressed, mice typically voluntarily head-fixed for greater than 10 minutes in one head-fixation session while maintaining a stable performance across different training days. Together with a participation rate of 100%, i.e., all mice in this study engaged in learning the task, our head-fixation mechanism potentially induced very little stress in mice, which helped to maintain the mice’s high motivation.

In this study we restricted the total amount of time one cage of mice had access to the automated training system, which equaled to 40 minutes times the number of mice in the cage, e.g., 2 hours for a cage of 3 mice, which is similar to the approach by Aoki et al. (2017), rather than giving mice access to the training system 24/7. However, we showed that the total head-fixation duration for each mouse in our system was close to 40 minutes, i.e., the same amount of time we assigned to each mouse on average, which means our system had a low idling time and was maximumly utilized by mice. We did not notice any dominating mice that occupied the training system for the majority of the time, and in fact all mice maintained good health across the training period (Figure 2Da), suggesting that all mice had the opportunity to collect enough water reward to maintain their body weight. Moreover, the restricted access to the training system potentially regulated the motivation level of the mice more consistently than allowing them access 24/7, which also mimicked the manual training protocol the most closely.

Furthermore, our study demonstrated that even with the restricted access the mice were able to perform around 300 trials per day. Compared to a previous study that trained mice on a tone detection task while giving mice 24/7 access (Francis *et al*., 2019), our paradigm resulted in a much higher daily trial number (∼300 vs. ∼70) and a higher hit rate (detection task: ∼60% vs. ∼6%). The restricted access opens the opportunity for a further increase in training throughput simply by automating the process where the animals are introduced and removed from the cage. In this scenario, each training setup can be used by multiple cages of animals per day and the system will operate with minimum idling time. Our automated head-fixation system thus has the potential to maximize the training capability of individual training setup, and it will be advantageous to combine such a setup with a high-end neural recording equipment, e.g., 2-photon microscope, that cannot be easily replicated at a low cost for each training setup. Such a combination can both maximize the training throughput as well as the utilization of the recording equipment.

In summary, we demonstrated that with our automated head-fixation training system, mice could achieve stable behavioral performance while maintaining a high level of participation and engagement. Our design has the potential to be integrated into other neural recording setup to maximize the throughput of both training and neural data collection.

## Acknowledgments

JL, and POK conceived study. JL and KM performed experiments. JL analyzed data. JL, POK wrote the manuscript. Supported by NIH R21MH116450 (POK), U19 NS107464 (POK).

